# The morphogenetic protein CotE drives exosporium formation by positioning CotY and ExsY during sporulation of *Bacillus cereus*

**DOI:** 10.1101/2020.12.11.422311

**Authors:** Armand Lablaine, Monica Serrano, Stéphanie Chamot, Isabelle Bornard, Frédéric Carlin, Adriano O. Henriques, Véronique Broussolle

## Abstract

The exosporium is the outermost spore layer of some *Bacillus* and *Clostridium* species and related organisms. It mediates interactions of spores with their environment, modulates spore adhesion and germination and could be implicated in pathogenesis. The exosporium is composed of a crystalline basal layer, formed mainly by the two cysteine-rich proteins CotY and ExsY, and surrounded by a glycoprotein hairy nap. The morphogenetic protein CotE is necessary for *Bacillus cereus* exosporium integrity, but how CotE directs exosporium assembly remains unknown. Here, we followed the localization of SNAP-tagged CotE, -CotY and -ExsY during *B. cereus* sporulation, using super-resolution fluorescence microscopy and evidenced interactions among these proteins. CotE, CotY and ExsY are present as complexes at all sporulation stages and follow a similar localization pattern during endospore formation that is reminiscent of the localization of *Bacillus subtilis* CotE. We show that *B. cereus* CotE drives the formation of one cap at both forespore poles by positioning CotY and then guides forespore encasement by ExsY, thereby promoting exosporium elongation. By these two actions, CotE ensures the formation of a complete exosporium. Importantly, we demonstrate that the assembly of the exosporium is not a unidirectional process as previously proposed but it is performed through the formation of two caps, as observed during *B. subtilis* coat morphogenesis. It appears that a general principle governs the assembly of the spore surface layers of *Bacillaceae*.

**IMPORTANCE:** Spores of *Bacillaceae* are enveloped in a glycoprotein outermost layer. In the *B. cereus* group, encompassing the *B. anthracis* and *B. cereus* pathogens, this layer is easily recognizable by a characteristic balloon-like appearance separated from the underlying coat by an interspace. In spite of its importance for the environmental interactions of spores, including those with host cells, the mechanism of assembly of the exosporium is poorly understood. We used super-resolution fluorescence microscopy to directly visualize formation of the exosporium during sporulation of *B. cereus* and we studied the localization and interactions of proteins essential for exosporium morphogenesis. We discovered that these proteins form a morphogenetic scaffold, before a complete exosporium or coat are detectable. We describe how the different proteins localize to the scaffold and how they subsequently assemble around the spore and we present a model for the assembly of the exosporium.

## INTRODUCTION

Bacterial endospores are one of the most resistant life forms (1). Endospores (hereinafter termed spores for simplicity) formed by members of the Firmicutes share a general morphological plan and are composed by several concentric layers. The core is the most internal structure and contains the bacterial chromosome. The core is surrounded by an “inner” membrane, then by a thin layer of peptidoglycan, the germ cell wall, and by an external and thicker layer of modified peptidoglycan, the cortex. The cortex, delimited by the outer forespore membrane (OFM), is enveloped in two main proteinaceous layers, the coat and the exosporium. While the coat is common to all species of spore-forming bacteria, the exosporium is only found in some *Bacillus* and *Clostridium* species and related organisms. In the *Bacillus cereus* group, encompassing the foodborne pathogen *B. cereus sensu stricto*, the etiologic agent of anthrax *B. anthracis* and the entomopathogenic species *B. thuringiensis*, the exosporium appears in transmission electron microscopy (TEM) as an irregular balloon-like structure, separated from the rest of the spore by the interspace, an electron transparent region of unknown composition. The exosporium is described as a thin “basal layer” with a paracrystalline structure surrounded by a “hairy nap” composed of different glycoproteins (2, 3). Whether the exosporium is somehow directly linked to the coat across the interspace remains unclear (4).

The exosporium directly contacts the environment, including host cells and the host immune system. It has a role in adhesion to abiotic surfaces and contributes to protection against predation and internalization by macrophages, as shown for *B. anthracis* spores (5). Despite its importance, assembly of the exosporium remains a poorly understood process. Electron microscopy reveals that its formation begins early in sporulation, at the onset of engulfment of the forespore by the mother cell, when the engulfing membranes start to curve, and before the coat or the cortex become recognizable (4, 6, 7). At this stage, the exosporium forms a single electron dense structure named the “cap” adjacent to the OFM at the mother cell proximal (MCP) forespore pole (4, 6, 7). After engulfment completion, the cap appears more markedly electrodense and becomes separated from the OFM by the interspace (4, 6, 7). The non-cap part of the exosporium is detected later, concomitantly with coat deposition (7). Assembly of the coat and exosporium appears to be interdependent, as recently shown in *B. anthracis* where formation of the cap part of the exosporium interferes with coat deposition at this site (7). Indeed, the coat first assembles along the longitudinal sides of the forespore, then at the mother cell distal (MCD) forespore pole and finally at the MCP forespore pole (7).

The exosporium is composed of at least 20 different proteins (5). Among them, ExsY, the main structural component of the exosporium basal layer, self-assembles into large 2-dimensional crystalline arrays (8). In absence of ExsY, only the cap part of the exosporium is assembled. CotY, sharing 90% amino acid sequence identity with ExsY, also self-assembles and probably forms the basal layer of the cap of *exsY* mutant spores (8–10). Importantly, spores of the *exsY cotY* double-mutant lack an exosporium (9). Both ExsY and CotY are cysteine-rich proteins, and cooperative disulphide bonds formation seems to play a role in their self-assembly (8). The current model of exosporium assembly proposes that CotY forms the cap possibly in association with ExsY (5, 8). From this cap, ExsY self-polymerizes to assemble unidirectionally towards the MCD pole, forming the remaining non-cap part of the basal layer. This structure offers a scaffold for assembly of additional exosporium proteins, such as BxpB required for the attachment of the collagen-like glycoproteins that form the hairy nap (11–17).

ExsY and CotY are orthologues of *B. subtilis* CotZ and CotY, which are important structural crust components (9, 18–20) and the self-organization of these cysteine-rich proteins plays an important role in formation of both layers (8, 21). Moreover, the morphogenetic protein CotE guides both outer coat and crust assembly in *B. subtilis* (18, 19, 22) and its *B. cereus* CotE orthologue is required for exosporium assembly (6, 23). These observations suggest that assembly mechanisms of crust and exosporium could be similar. The assembly of the proteins forming the different layers of *B. subtilis* coat is a sequential process divided in two steps. First, a group of early-synthesized proteins, mainly morphogenetic proteins, forms an organizational center on the MCP forespore pole, named the cap, which is dependent on the morphogenetic ATPase SpoIVA (20, 24). CotE and CotZ are part of this cap (20). In a second step, the different coat proteins encapsulate the circumference of the spore from the single cap, a process named encasement that requires SpoVID (25). Encasement by CotE depends on a direct interaction with SpoVID and involves formation of a second cap at the MCD following engulfment completion, at the site of membrane fission (20, 25, 26). Other proteins, designated as kinetic class II proteins (20), share this sequence of localization and encasement.

How the exosporium proteins localize during the course of sporulation, which proteins dictate the different steps of exosporium assembly and on which interactions the process relies are unknown in the *B. cereus* group. In particular, how CotE contributes to the assembly of the exosporium is unknown. Here, we study the localization and the dependencies between CotY, ExsY and CotE during *B. cereus* sporulation using super-resolution fluorescence microscopy. We show that CotY, ExsY and CotE exhibit the same pattern of assembly throughout sporulation that is reminiscent of the localization of the kinetic class II coat proteins of *B. subtilis* to which CotE belongs. Importantly, our results also reveal that CotE forms complexes with CotY and ExsY during sporulation and that CotE directly interacts with these proteins. CotE drives CotY and ExsY assembly via direct and indirect interactions, allowing the formation of two cap structures and then elongation of the exosporium from these caps. These two crucial steps ensure the progressive assembly of the exosporium around the spore. These features are highy similar to those observed during coat layers formation in *B. subtilis*, suggesting that the principle governing assembly of the spore surface layers is conserved among distant members of the *Bacillus* group.

## RESULTS

### CotY, ExsY and CotE follow the same assembly dynamic during sporulation

To determine the role of CotE, together with CotY and ExsY in the formation of *B. cereus* exosporium, we first monitored the localization of these three proteins during sporulation using super-resolution structured illumination microscopy (SR-SIM) (27). We used translational fusions to the SNAP-tag, which becomes fluorescent (red signal on overlay images) upon covalently binding of the TMR-Star substrate (TMR) (28–31). The role of CotE in *B. cereus* was mostly studied in ATCC 14579 strain and previous work showed that more CotE was extracted in spores formed at 20°C than at 37°C (23). For this reason, sporulation of ATCC 14579 producing CotY-, ExsY-, or CotE-SNAP fusions (SNAP at the C-terminal end of the proteins) or SNAP-CotE (SNAP at the N-terminal end of CotE) was performed at 20°C. We checked with conventional fluorescence microscopy that CotY-SNAP and ExsY-SNAP followed identical patterns of localization during sporulation at 20° C and 37° C (data not shown). Culture samples were collected throughout sporulation, labeled with TMR and the different patterns of SNAP-fusion localization were scored with respect to the stages of sporulation identified using the Mito Tracker Green dye (MTG, green fluorescent signal on overlay images) (32).

We first detected CotY-SNAP as a discrete fluorescence signal on curved septa at the onset of engulfment (Fig. 1, pattern *a*, white arrow) in 13% and 2 % of the sporangia scored after hour 20 and hour 24 of sporulation, respectively (Fig. S1C). This signal then extended along the engulfing membranes and became cap-shaped representing 33% and 7% of the sporangia after hour 20 and 24 of sporulation, respectively (Fig. 1 and S1C; pattern *b*). Importantly, the cap pattern of CotY-SNAP persisted until the end of engulfment, representing 33% and 25% of the sporangia at hour 20 and 24 of sporulation, respectively (Fig. 1 and S1C, pattern *c*). At hour 24, we distinguished different patterns of CotY-SNAP localization in sporangia having completed engulfment, *i*) a cap-shaped red fluorescent signal at the MCP pole in 25% of the sporangia (Fig. 1 and S1C, pattern *c*); *ii*) caps at both forespore poles in 10% of the sporangia (pattern *d*); *iii*) a red fluorescent signal covering three quarters of the forespore in 26% of the sporangia (pattern *e*) and *iv*) a ring of red fluorescence in 30% of the sporangia (pattern *f*). Remarkably, in fully engulfed sporangia, a layer labeled by MTG was detected at the MCP forespore pole and separated from the forespore membrane (Fig. 1, pink arrows, patterns *d, e* and *f*). Strikingly, the CotY-SNAP signal appeared to migrate from the inner forespore membrane (Fig. 1, pattern *c*) to this nascent layer (patterns *d, e* and *f*). Later, this layer exhibited the recognizable balloon-like structure of the exosporium and completely surrounded the spore, showing that MTG also labeled the exosporium (Fig. 1, pink arrows, patterns *g* and *h*). Hence, this layer, when detected at the MCP forespore pole in sporangia that had completed engulfment, likely corresponds to the cap (patterns *d, e* and *f*). From hour 48, only sporangia of phase bright spores and free spores were observed by phase contrast microscopy (data not shown) and the CotY-SNAP signal clearly superimposed onto the exosporium basal layer (Fig. 1, pattern *g*). Then, the signal was more diffused in free spores (Fig. 1, pattern *h*). Similar localization patterns were observed for ExsY-SNAP and SNAP-CotE (Fig. S1A, B and C). The dynamics of CotE-SNAP assembly was globally similar, with slight differences in sporangia that had completed the engulfment process (Fig. S2 and Supplemental text).

**FIG 1.**
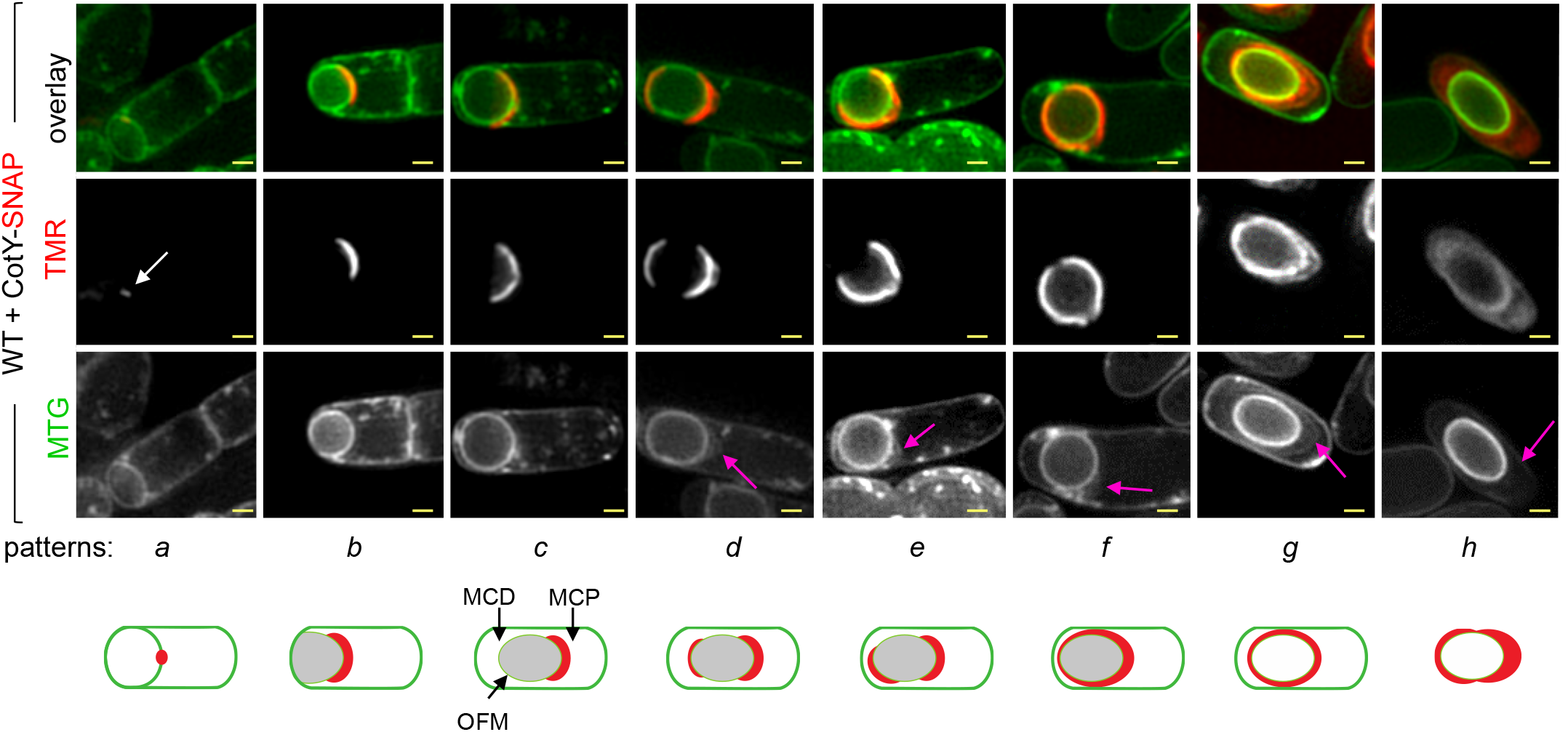
Stages of CotY-SNAP localization during sporulation. Sporulating cells of *Bacillus cereus* producing CotY-SNAP were labeled with the SNAP substrate TMR-Star and with the membrane dye MTG and imaged by SR-SIM microscopy. Images of a representative cell of each identified pattern of localization at different stages of sporulation are shown with a schematic representation at the bottom of the panels (patterns *a* to *h*). White arrow points to the weak signal of CotY-SNAP detected at curved septa ; pink arrows point to the exosporium at different stages of its formation; black arrows point to the mother cell proximal (MCP) and distal (MCD) forespore poles and to the outer forespore membranes (OFM). Scale bars, 0.5 µm. The stages of ExsY-SNAP, SNAP-CotE and CotE-SNAP localization during sporulation are shown on Figure S1A-B and S2A, respectively.

Taken together, these results show that CotY, ExsY and CotE share a common pattern of localization during sporulation, behaving as cap proteins at the MCP forespore pole during engulfment, then after engulfment completion forming a second cap at the MCD spore pole and finally completely encasing the forespore. Thus, CotY and ExsY do not seem to be restricted to the cap or the non-cap regions of the *B. cereus* exosporium, as previously proposed (5, 12, 33). We also found that CotY, ExsY and CotE already encased the forespore in most of the sporangia scored at hour 24 (Fig. 1 and S1, pattern *f*). Importantly, the coat were not detected and only the cap region of the exosporium was visible by TEM at this time (Fig. S3B). Hence, CotE, CotY and ExsY are present at the surface of the non-cap region of the forespore at a time the exosporium is not yet detected by TEM in this region of the forespore.

### CotE is required for the localization of CotY and ExsY as two caps

We used SR-SIM to analyze the localization of CotY and ExsY in *cotE* sporangia and in mature spores. Neither caps formation nor encasement by CotY-SNAP or by ExsY-SNAP was observed in the absence of CotE (Fig. 2A and B, panels *a* to *f*). Remarkably, in the absence of CotE, CotY-SNAP accumulates as dots at the forespore poles, while ExsY-SNAP formed large aggregates in the mother cell, often close to the poles (Fig. 2A and B respectively, purple arrows in panel *f*).

**FIG 2.**
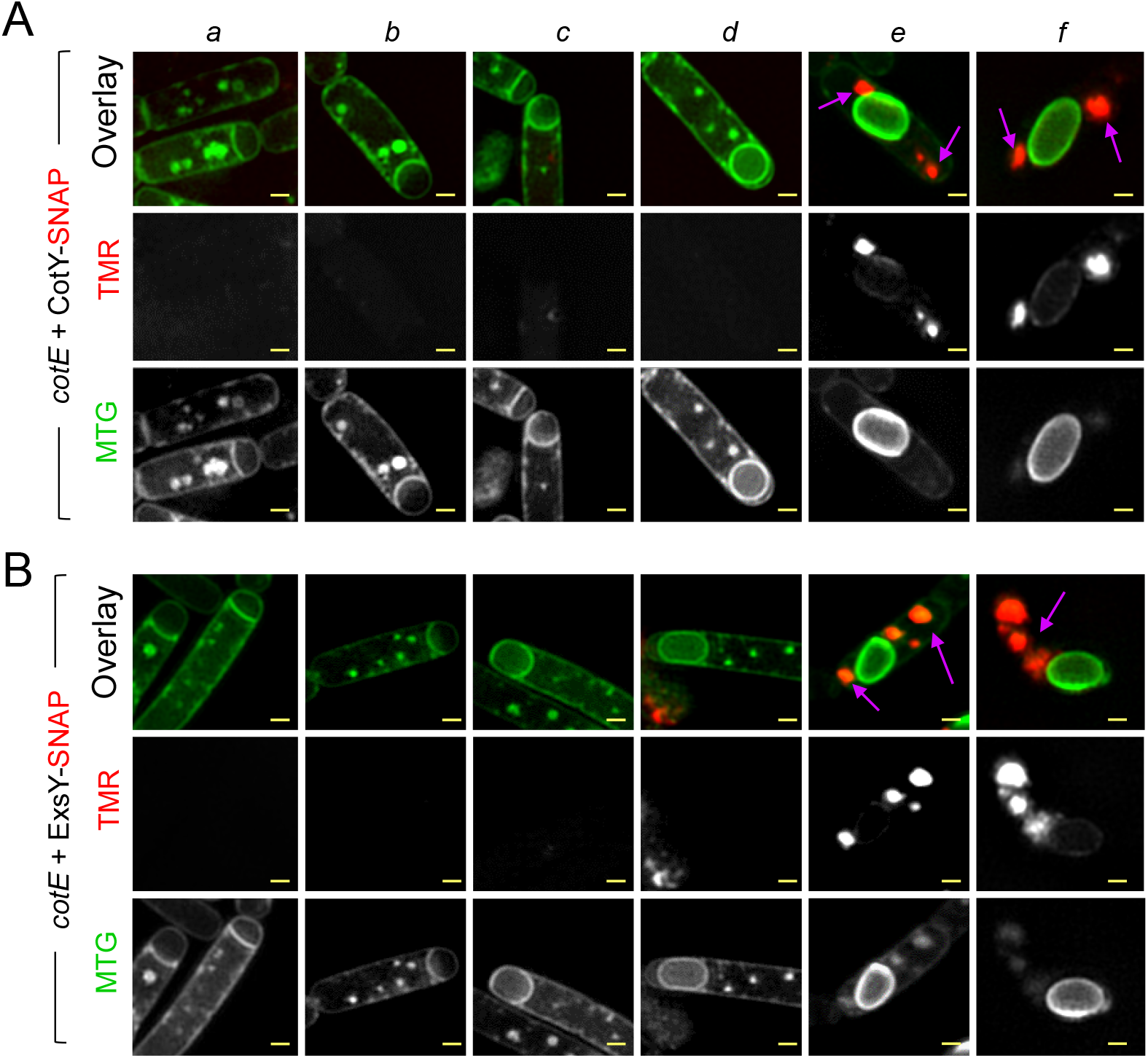
CotE is required for cap formation by allowing CotY and ExsY to localize as caps. Samples were collected from cultures of a *cotE* mutant producing CotY-SNAP (A) or ExsY-SNAP (B) during sporulation and the cells imaged by SR-SIM following staining with TMR-Star and MTG. Representative cells of the different localization patterns observed are shown (panels *a* to *f*). Purple arrows point to the signal from CotY-SNAP or ExsY-SNAP (panels *e* and *f*). In the *cotE* mutant, CotY-SNAP and ExsY-SNAP never formed the cap nor the following patterns identified in WT (Fig. 1 and S1A) but formed large patches or aggregates in the mother cell cytoplasm. Scale bars, 0.5 µm.

As CotY and ExsY cap-shaped signals were never seen in *cotE* mutant sporangia, we reasoned that the cap could be absent in this strain. With TEM, we confirmed the absence of cap formation and we found that in the absence of CotE, deposition of the coat follows a modified pathway (Fig. S3E-G and Supplemental text). Both fluorescence microscopy and TEM show that CotE is required to form the cap of the exosporium by allowing CotY and ExsY localization first as a cap at the MCP forespore pole and then as a second cap at the MCD forespore pole. This localization pattern and/or the presence of CotE appears important to guide the subsequent encasement of the forespore by CotY and ExsY.

### ExsY is required for encasement by CotY but not for encasement by CotE

CotY, ExsY and CotE appeared to encase the forespore early, in sporangia presenting only the cap region of the exosporium. Thus, we wondered whether the assembly of CotY and CotE is affected by the absence of ExsY, as only the cap region of the exosporium is assembled in *exsY* endospores (10). To test whether the localization of CotY and CotE requires ExsY, we used an *exsY* mutant previously obtained in ATCC 10876 strain (9). The amino acid sequence of *B. cereus* ATCC 10876 and ATCC 14579 CotE are 100% identical and 92.9% identical for CotY. Moreover, we showed that the CotY-SNAP construction from the *B. cereus* ATCC 14579 CotY sequence complements the *cotY* mutation in ATCC 10876 (Fig. S4C). For this strain, however, and for reasons we do not presently understand, when sporulation is induced at 20°C, engulfment is stopped in a fraction of the cells (not shown). We therefore induced sporulation at 37°C: in *exsY* mutant sporangia, CotY-SNAP localized as a cap at the onset of engulfment (Fig. 3A, pattern *b*) and until the end of engulfment (pattern *c*); together these patterns represented 64% of the cells observed at hour 12. In engulfed sporangia at hour 12, CotY-SNAP formed a second smaller cap at the MCD forespore pole (pattern *d*) in 13% of the cells and localized as a cap at the MCP pole and as a weak dot on the MCD pole (pattern *j*) in 19% of the cells. Importantly, the localization of CotY-SNAP did not change over time, suggesting a blockage of CotY assembly. Patterns *c, d* and *j* were observed in phase bright endospores and spores (Fig 3A, patterns noticed *c*\* to *j*\*). From hour 14, an inner signal was detected in phase bright sporangia (Fig. 3A, cyan arrow), likely due to unspecific binding of the TMR to the forespore, as observed in a strain without a SNAP fusion (data not shown). In free spores observed at hour 34, CotY-SNAP never formed the fluorescent ring fully encasing WT spores. These results show that ExsY is required for CotY encasement but not for its localization as two caps. Considering that the exosporium cap is made of CotY in *exsY* sporangia, these results suggest that CotY could form the basal layer of the cap independently of ExsY but relying on CotE.

**FIG 3.**
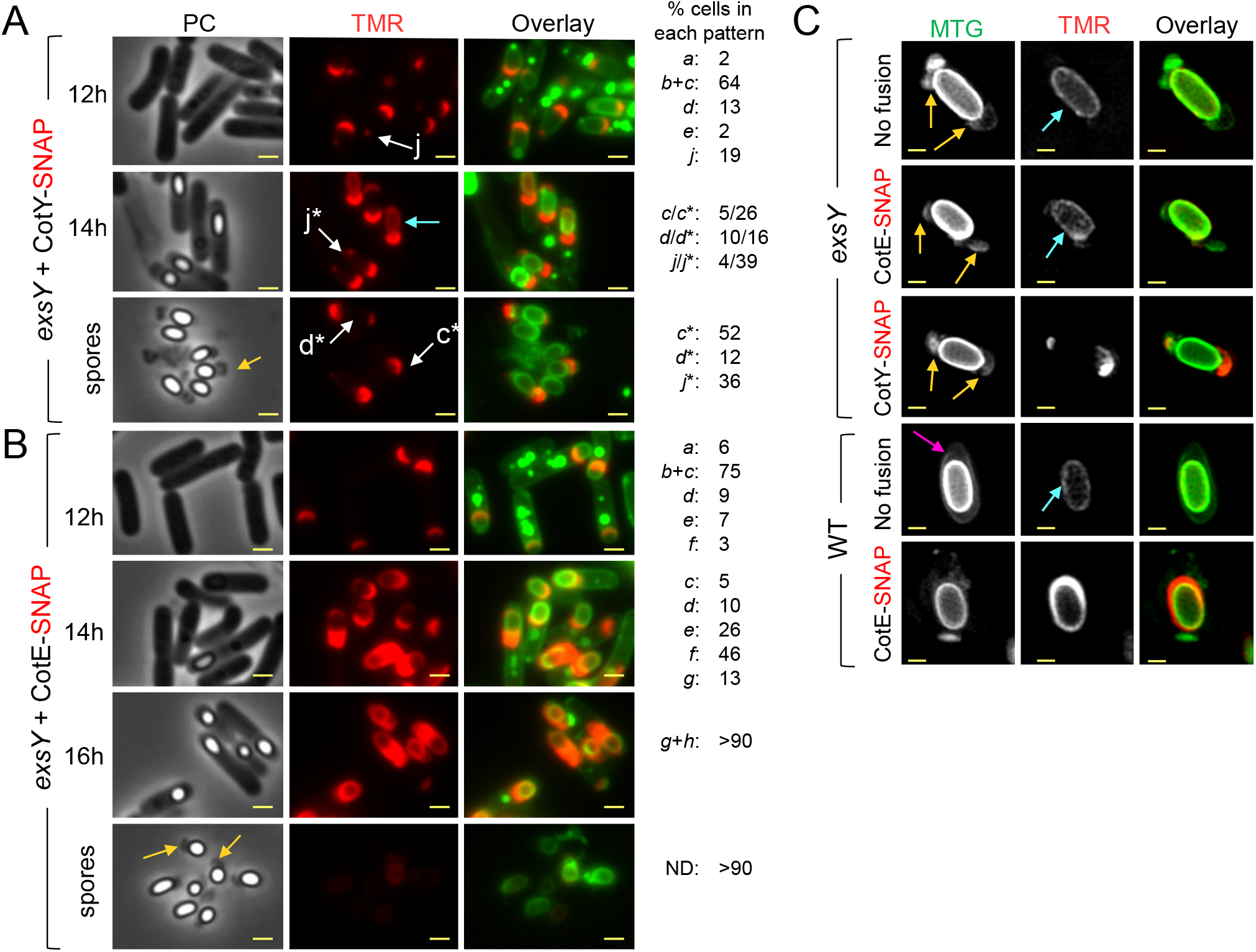
ExsY is required for encasement by CotY but not by CotE. Samples were withdrawn at the indicated times from sporulating cultures at 37°C of an *exsY* mutant producing CotY-SNAP (A) or CotE-SNAP (B). The cells were labeled with TMR-Star and MTG and imaged by phase contrast and fluorescence microscopy. The localization patterns *a* to *g* are identical to the *a*-*g* patterns observed in WT cells in Fig.1. Patterns *c*\*, *d*\*, *j*\* (white arrows) correspond to patterns *c, d, j,* observed in sporangia of phase bright spores or in free spores. The numbers on the right side of the panels show the percentage of cells with each of the indicated patterns, relative to the total number of sporulating cells expressing the different SNAP-fusions. ND; no signal detected. (C) Spores of indicated strains were stained with MTG and TMR-STAR and imaged by SR-SIM microscopy. The strains were ATCC 10876 (WT) and a derivative producing CotE-SNAP; a congenic *exsY* mutant and derivatives producing CotE-SNAP or CotY-SNAP. Yellow arrows point to an exosporium blocked at the two caps stage. The pink arrow points to a complete exosporium, cyan arrows point to a non-specific TMR signal. Scale bar in panels A and B, 1 µm and in C, 0.5 µm.

The sequence of CotE-SNAP and SNAP-CotE localization in *exsY* sporangia (Fig 3B and S4A) were both similar to the ones in the WT (Fig. S1B and S2A). Interestingly, no signal from CotE-SNAP and SNAP-CotE was observed in the cap region of *exsY* spores after hour 34, despite a visible cap structure in phase contrast images (Fig. 3B and S4A, yellow arrows). These observations suggest that CotE is not present or not detectable in *exsY* spores, although it correctly assembles during sporulation. A faint inner red TMR signal was observed, possibly corresponding to unspecific TMR binding to phase bright forespores (Fig. 3B). To check this, we used SR-SIM to analyze *exsY* spores of strains producing CotY-SNAP, CotE-SNAP or from a strain bearing no SNAP-fusion. As suggested by conventional fluorescence microscopy (Fig. 3B), we only detected an internal faint TMR signal in *exsY* spores with CotE-SNAP (Fig. 3C, cyan arrow), also seen in WT and *exsY* spores without SNAP-fusion. In spores of *exsY* producing either CotY-SNAP or CotE-SNAP and presenting a two caps MTG signal (Fig. 3C, red arrows), we clearly observed fluorescence from CotY-SNAP-TMR (Fig. 3C) while no signal was observed for CotE-SNAP (Fig. 3C). This suggests that CotY was present in the cap of the *exsY* mutant, while CotE was not (Fig. 3C). These results confirm the absence of a detectable signal from CotE-SNAP in spores lacking ExsY, even in the cap region and that a second cap is formed by CotY during exosporium formation. We used immunoblotting to test for the presence or absence of CotE in *exsY* and in *cotY* spores (Fig. S4B). In line with microscopic analysis, CotE was not detected in *exsY* spores; but more surprisingly, it was not detected in *cotY* spores either (Fig. S4B).

Altogether, these results show that CotE assembly is independent of ExsY and of CotY, as ExsY is required for CotY assembly. However, the absence of ExsY or CotY possibly affects the association of CotE with the released spores. In addition, while CotE completely encased *exsY* forespores, CotY assembly was blocked at the two caps stage. Thus, encasement by CotE is not sufficient to direct encasement by CotY, which additionally requires ExsY.

### CotE, CotY and ExsY form a complex *in vivo* during sporulation and interact *in vitro*

As CotE is required for the correct localization of CotY and ExsY during sporulation, we wondered whether CotE could form complexes with CotY and ExsY during spore formation. We used the ability of SNAP-tag to covalently bind SNAP-capture agarose-beads (34) to follow *in vivo* complex formation between CotE, CotY-SNAP and ExsY-SNAP. Pull-down assays were performed on extracts prepared from sporangia early after engulfment completion (hour 20), sporangia at the end of the engulfment process (hour 24), phase bright endospores and spores (hour 48 and hour 72, respectively). We also performed pull-down assays on extracts from purified spores collected at hour 72. In line with the microscopy observations, we observed accumulation of CotE (Fig. 4A, extracts panels), CotY-SNAP and ExsY-SNAP (Fig. S5A, extracts panels) from hour 20 to 72 of sporulation. CotE was extracted as a species of about 20 kDa corresponding to the expected size of the monomer, an abundant species of about 40 kDa, possibly corresponding to a dimer and a less abundant multimeric form of about 75 kDa. We also detected species of intermediate apparent molecular weight, which may correspond to cleaved forms of CotE. SNAP fusion proteins were detected in the supernatant, after incubation of extracts collected from hour 48 with SNAP-beads (Fig. S5A). All SNAP fusions were detected with anti-SNAP antibodies after the pull-down, and since the SNAP moiety covalently binds to SNAP-capture beads, this suggests that the SNAP fusions could self-interact (Fig S5A, pull-down panels). We found that CotY-SNAP and ExsY-SNAP pulled-down the different forms of CotE extracted from sporulating cells and from purified spores (Fig 4A, pull-down panels). Notably, the high molecular weight CotE species seems to be preferentially enriched in the pull-down compared to the extracts (Fig. 4A, three red asterisks). In contrast, a non-specific species (green asterisk in Fig. 4A) detected from hour 0 (*i*.*e*. vegetative cells) was present in extract and flow-through fractions but absent in the pull-down. The SNAP alone, produced under the control of the *cotE* promoter (P*cotE*-SNAP fusion) did not retain CotE (Fig 4A, pull-down panels). These results show that CotE formed complexes with CotY and with ExsY during sporulation and in mature spores of *B. cereus*. We also performed pull-down SNAP assays on extracts of sporulating *exsY* cells harboring CotY-SNAP, in which CotY assembly was blocked at the two caps stage (Fig 3A and S6B). The results indicate that CotE and CotY are present in complexes in the two caps and that the formation of these complexes is independent of the presence of ExsY (Fig. S6A and see Supplemental material).

**FIG 4.**
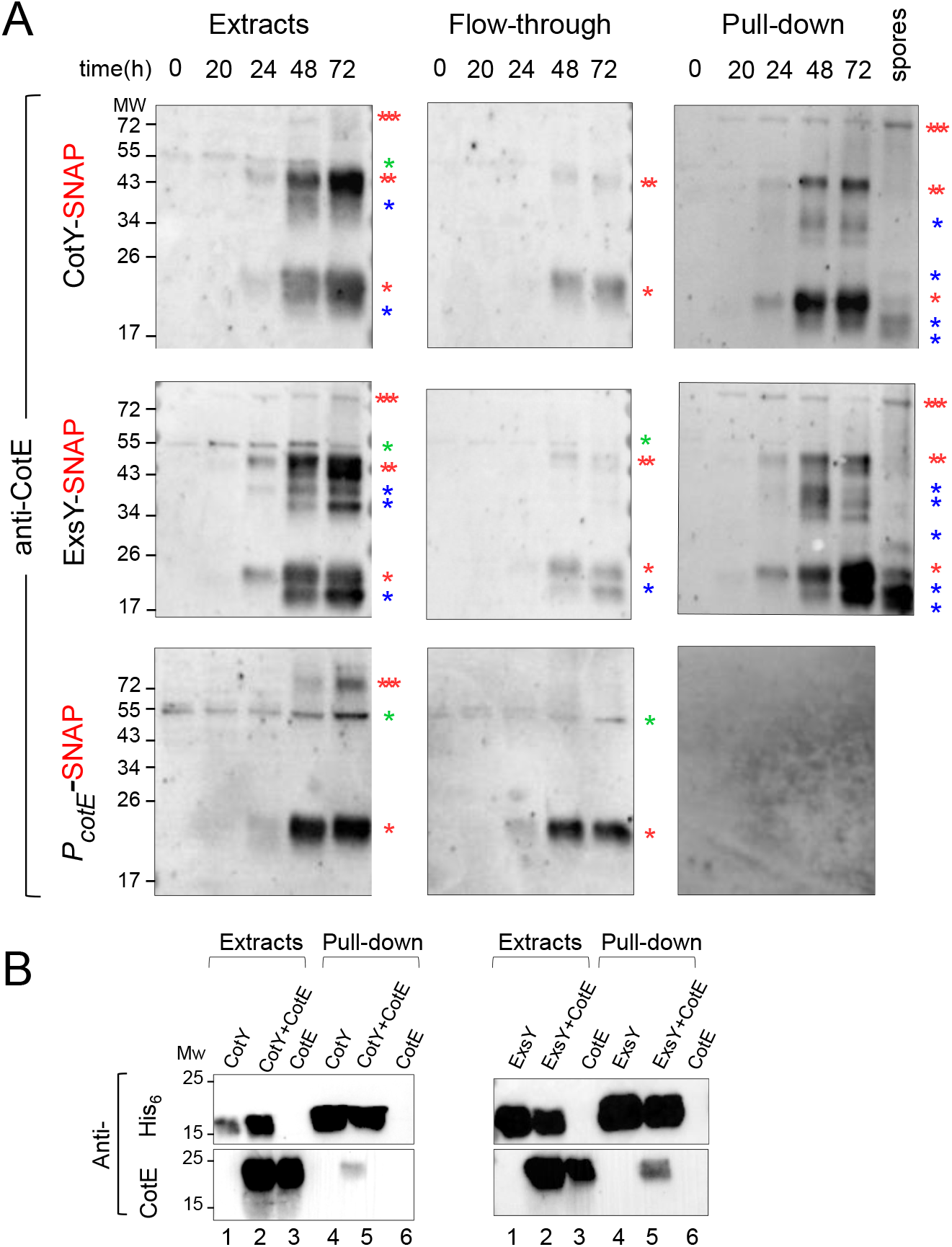
CotE, CotY and ExsY interact *in vivo* and *in vitro*. Samples were collected from sporulating cultures of *B. cereus* strains producing various SNAP fusions at the indicated times. Whole cell extracts were prepared and submitted to pull-down assays with a SNAP-capture resin. Whole cell extracts, flow-through and bound proteins were resolved by SDS-PAGE and subjected to immunoblot analyses with anti-CotE antibodies (A). A lane corresponding to a pull-down realized on proteins extracted from purified CotY-SNAP and ExsY-SNAP spores was added. One red asterisk indicates a monomer of CotE, two red asterisks a potential dimer of CotE and three red asterisks indicate multimers; blue asterisks indicate possible proteolytic products of CotE. A non-specific signal in the extracts from vegetative cells (hour 0) is indicated by a green asterisk. (B) Heterologous co-expression pull-down assays. The indicated protein from *E. coli* BL21 (DE3) cell lysates were submitted to pull-down analysis. Presence/absence of the proteins was tested by immunoblot analysis with an anti-His_6_ antibody (upper panels) or with an anti-CotE (lower panels for co-production of CotY-CotE and ExsY-CotE). Lanes 1, 2, 3 illustrate synthesis of the various proteins present in the different cell extracts. Lines 4, 5, 6 show the elution analysis after pull-down assays. In A and B, the position of molecular weight markers (MW, in kDa) is shown on the left side of the panels.

To determine if CotE interacts directly with CotY and/or ExsY, the proteins were co-produced in *E. coli* and tested for pairwise interactions (35). Co-expression using pETDuet system allows formation of interactions in the cell and purification of protein complex (35). Notably, organized supramolecular structures formed by *B. subtilis* coat proteins were purified with this approach (21). To determine possible interactions, we co-expressed CotE with His_6_-CotY or His_6_-ExsY. We observed that His_6_-CotY, His_6_-ExsY and CotE accumulated in *E. coli* and were detected as abundant species migrating at the expected size of the monomer, i.e. 18.2 kDa for His_6_-CotY, 17.5 kDa for His_6_-ExsY (Fig 4B; lines 1 and 2 in upper panels) and 20.3 kDa for the native CotE (Fig 4B; lines 2 and 3 in bottom panels). In line with previous results (8), His_6_-CotY or His_6_-ExsY were detected as dimers and as high molecular complexes despite the addition of urea, showing the difficulty to solubilize these complexes (not shown). His_6_-CotY and His_6_-ExsY bound and were eluted from the Ni^2+^-beads, mainly as monomers (Fig 4B; lines 4 and 5 in upper panels) and CotE was not retained by the Ni^2+^-beads when produced alone (Fig 4B; line 6, bottom panels). When co-produced with His_6_-CotY or His_6_-ExsY, CotE was eluted together with His_6_-CotY and His_6_-ExsY (Fig 4B; lines 5) showing that CotE can directly interact with both CotY and ExsY.

## DISCUSSION

Here, we examined the assembly of the exosporium in *B. cereus* focusing on CotY and ExsY, and their relationship with CotE. We show that CotY, ExsY and CotE share a common pattern of localization and are detected as complexes throughout sporulation and CotE can directly interact with CotY and ExsY. CotY, ExsY and CotE co-localize at the MCP forespore pole during engulfment and the combination of fluorescence microscopy and TEM reveals that CotE is required to form the cap region of the exosporium (Fig. 5A*i*). Following engulfment completion, the three proteins co-localize as a second cap at the MCD forespore spore pole (Fig. 5A*ii*). CotE is required for the localization of both ExsY and CotY at both caps. In the absence of CotE, CotY-SNAP forms dots at the forespore poles and ExsY-SNAP becomes dispersed throughout the mother cell cytoplasm. Finally, from the two caps, the three proteins encase the forespore (Fig. 5A*iii-iv*). Thus, the localization of CotY and ExsY is not confined to the cap or to the non-cap regions of the *B. cereus* exosporium, respectively, as previously thought (5, 12, 33).

**Fig 5.**
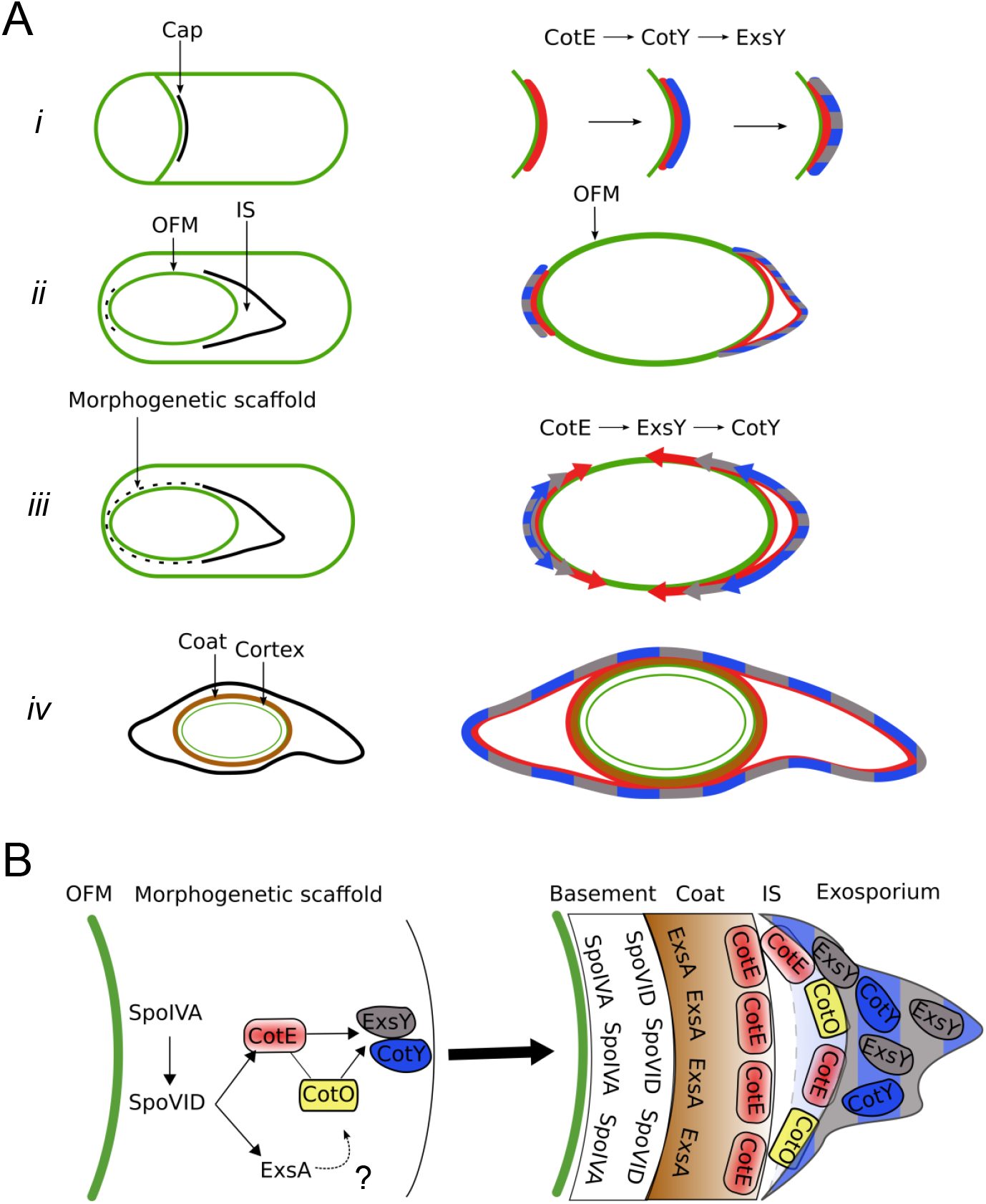
Successive localization, interactions and interdependence among CotE, CotY and ExsY during exosporium formation. (A) *i* to *iv* represent the assembly of the indicated spore structures as observed by TEM. Dotted lines show the new structures found in the present study. The diagrams on the right side show the temporal sequence of localization of CotE, CotY and ExsY inferred from our results. (*i*) CotE (red) forms a cap in the septal region at the onset of engulfment and recruits CotY (blue) which in turn recruits ExsY (grey). Once positioned, CotY and ExsY form the basal layer of the cap (mixed grey and blue area). The cap remains unchanged until engulfment completion. (*ii*) After engulfment completion, a second smaller cap (dotted line), is formed at the MCD forespore pole through the localization of CotE, first, then CotY and finally ExsY. At the same time the MCP forespore cap separates from the OFM and the interspace (IS) begins to be formed. (*iii*) CotE progressively encases the spore (red arrows) guiding the simultaneous encasement of ExsY (grey arrows) that in turn is required for encasement by CotY (blue arrows). CotE, CotY and ExsY thus localize around the forespore as a morphogenetic scaffold. (*iv*) After coat formation, CotE, CotY and ExsY are found around the entire spore, and the exosporium forms a continuous layer. (B) The formation of the morphogenetic scaffold and its maturation in *B. cereus*. Based on their roles in *B. subtilis*, SpoIVA and SpoVID homologs are good candidates to direct recruitment and encasement, respectively of CotE and by CotE and CotE-controlled proteins. CotO, possibly through its interaction with CotE, may participate in the recruitment of CotY and/or ExsY. ExsA, a SafA homologue that controls *B. cereus* coat protein deposition is also present in the morphogenetic scaffold. Whether ExsA is implicated in the recruitment of CotE and/or CotE-controlled proteins in the morphogenetic scaffold is unknown. After sporulation completion, the proteins of morphogenetic scaffold are positioned in the different layers of the mature spore.

Certain aspects of this pathway deserve special attention. We were able to distinguish with SR-SIM the cap part of the exosporium at the time of its separation from the OFM by the formation of the interspace, just after engulfment completion. At this stage in morphogenesis, we show that CotY, ExsY and CotE migrate from the OFM to form the visible basal layer of the cap region of the exosporium and to start encasement of the forespore (Fig. 5A*ii-iii*). How the separation of CotY, ExsY and CotE to form the cap is brought about is unclear. With a CotE-SNAP fusion, we noticed the cap to be closer to the OFM, suggesting that the SNAP interferes with a function of the C-terminal region of CotE in the separation of the cap from the OFM and more generally, that CotE may be implicated in interspace formation. However, this question, often raised (5, 22) has to be further investigated.

The second cap that forms at the MCD forespore pole after engulfment completion (Fig. 5A*i*, dotted line), is smaller than the cap at the MCP pole, and was detected only in *exsY* sporangia where exosporium assembly is blocked, but not in WT sporangia. This suggests that the second cap is a transient structure, compared to the cap at the MCP forespore pole, which persists throughout engulfment. Importantly, this result demonstrates that the assembly of the exosporium is not a unidirectional process, starting from the MCP and progressing towards the MCD pole as previously proposed (5, 7, 8, 10). Rather, it involves the formation of two organizational centers, the caps, at the forespore poles (Fig. 5A*ii*). Both caps are dependent on CotE, can be formed by CotY independently of ExsY and relies on a CotE-CotY interaction (Fig. 5A*ii*).

The last events in the assembly of CotY, ExsY and CotE around the forespore follow a particular dynamics, in that a three-quarter of a circle localization pattern (pattern *e*) is detected. While CotY-SNAP localized as two caps (pattern *d*) in *exsY* sporangia, pattern *e* was not observed, showing that the localization as a three-quarter circle occurred after the localization as two caps, and that this localization of CotY relies on ExsY. Also, since the three-quarter circle localization of CotE is independent of ExsY, it seems that CotE imposes this localization to ExsY and thus to CotY (Fig. 5A*iii*). Recently, Boone *et al* reported that the assembly of the exosporium in a *cotY* mutant, and therefore assembly of ExsY, initiates at random locations (7). Based on our results, we propose that the encasement of the forespore by ExsY in a *cotY* mutant is allowed by a CotE-ExsY interaction, as CotE asymmetric encasement appears independent of CotY encasement.

Importantly, encasement by CotY, ExsY and CotE was completed before the appearance of the non-cap part of the exosporium or coat formation, as observed by TEM. We propose that these proteins form a morphogenetic scaffold at the surface of the non-cap part of the forespore. Previous studies suggested that a region named the spore-free sacculus or “sac”, is implicated in exosporium assembly, allowing attachment of the cap in *exsY* sporangia as well as assembly of ExsY (8, 10, 22). The morphogenetic scaffold we describe here seems to correspond to this “sac”. If so, sac formation largely relies on the interactions of CotE with CotY and ExsY involved in the formation of the caps and exosporium elongation.

The assembly pathway of CotY, ExsY and CotE is reminiscent of that of class II coat proteins in *B. subtilis*, which form a MCP cap and a MCD cap following engulfment completion when encasement starts. There are, however, differences to be highlighted. *B. subtilis* CotZ is a class III protein, *i*.*e*, it begins encasement only when the forespore turns phase dark. However, in *B. subtilis*, but not in *B. cereus*, transcription of *cotY* and *cotZ* is under the control of GerE, which could explain the different behavior as encasement also depends on waves of gene expression (20, 36). In addition, no asymmetric encasement has been reported in *B. subtilis*. Finally, in the absence of CotZ, *B. subtilis* CotY fails to form a second cap (37).

Nevertheless, the assembly of the spore surface layers may follow common principles across species. The *B. subtilis* coat initially assembles as a scaffold composed of proteins present in the four layers of the mature coat, including SpoIVA, SpoVID, SafA, CotE and CotZ. SpoIVA recruits proteins to the spore surface, SpoVID controls encasement, SafA controls assembly of the inner coat, CotE directs assembly of the outer coat and together, CotE and CotZ govern assembly of the crust (18-20). SpoIVA and SpoVID are conserved in *B. cereus* where a *spoIVA* mutation leads to important defects of exosporium and coat assembly, similarly to *B. subtilis* (6, 22). It is likely that SpoIVA and SpoVID have similar roles in *B. cereus* (Fig. 5B). Recently, it was shown that the GerP coat proteins did not localize properly in a *B. cereus exsA* mutant but localized normally in a *cotE* mutant, suggesting a separate control of coat and exosporium assembly (38). ExsA, a paralog of *B. subtilis* SafA, is transcribed just after initiation of the sporulation and while an *exsA* mutation impairs exosporium assembly, this is likely an indirect effect due to the misassembly of the internal coat (39). Thus, ExsA may also be present in the morphogenetic scaffold of *B. cereus* that we propose here (Fig. 5B) and may have a role equivalent to that of SafA in *B. subtilis* (40). Also recently, in *B. anthracis*, CotO was shown to share similar characteristics with CotY and ExsY. CotO is an orthologue of a class II morphogenetic protein implicated in the encasement of the spore by crust proteins in *B. subtilis* (20, 37). *B. anthracis* CotO interacts with CotE, is dependent of CotE for its localization and shows a potential class II localization kinetics (7). Thus, it is tempting to propose that CotO is part of the morphogenetic scaffold of *B. cereus* together with CotE, CotY and ExsY (Fig. 5B). Interestingly, CotO is required for assembly of the exosporium and *cotO* mutation leads to a phenotype similar to a *cotE* mutation (7), suggesting that CotE and CotO work in concert, possibly by stabilizing the complexes formed by CotE, CotY and ExsY that we describe here.

Formation of the coat scaffold of *B. subtilis* and the dynamics of the coat/crust proteins is driven by self-polymerization and by direct interactions (21, 24). Similarly, it has been shown that CotY and ExsY could self-polymerize and directly interact in *B. cereus* (8). We found that ExsY polymerization seems not dependent of CotE and is not required for CotE assembly around the spore but appears necessary for the assembly of CotY. The crosslinking, via formation of disulfide bonds between ExsY monomers and with CotY, could be a driver of exosporium formation. The resulting ExsY/CotY expansion around the developing spore is driven by a sublayer of CotE which, in turn, may rely on a SpoIVA/SpoVID plateform. Thus, CotE and the CotE-controlled proteins, CotO, CotY and ExsY, are recruited in the *B. cereus* morphogenetic scaffold (Fig. 5B).

The incorporation of a second group of late synthesized proteins, such as CotB, ExsB, ExsK, BxpB, BetA, or BclA, could allow the maturation of the coat and exosporium structures (3, 15, 16, 36, 41–44). Most of these proteins are unique to the *B. cereus* group and are likely responsible for the distinctive properties of the spore surface layers (22). In any case, regardless of the important structural differences observed between the *B. subtilis* and *B. cereus* proteinaceous layers, a scaffold made of morphogenetic proteins fulfills the same role by guiding the localization of other components through direct protein-protein interactions.

## MATERIALS AND METHODS

### Bacterial strains and growth conditions

Bacterial strains and plasmids used in the present study are listed in Table S1. LB broth with orbital shaking (200 rpm) or LB agar plates were used for routine growth of *B. cereus* and *E. coli* at 37°C. When needed, liquid cultures or plates were supplemented with antibiotics at the following concentrations: ampicillin (Amp) at 100 µg.ml-1 for *E. coli* cultures, spectinomycin (Spc) at 275 µg.ml-1, and erythromycin (Erm) at 5 µg.ml-1 for *B. cereus* cultures.

### Other general methods

The construction of SNAP fusions and plasmids for the co-production of proteins in *E. coli* as well as the methods used for protein co-production, pull-down assays and immunoblotting analysis are described in the Supplemental material.

### Sporulation kinetics

Sporulation was induced in liquid SMB medium at either 20°C or 37°C with orbital shaking at 180 rpm, as previously described (23). Cells were harvested at the desired time by centrifugation for 10 min at 10,000 x g before microscopy or immunoblotting analysis. When necessary, spores were collected after 72 h of incubation and purified by successive centrifugations and wash with cold water as described (23). For immunoblot analysis, *B. cereus* spores were also produced at 20°C on modified fortified nutrient agar (mFNA) (23).

### SNAP labelling and fluorescence microscopy

Samples (5 to 10 ml) were withdrawn from cultures in SMB at selected times during sporulation. Cells were collected by centrifugation (10,000 x g for 10 min), suspended in 200 µL of phosphate saline buffer (PBS) and labeled with TMR-Star (New England Biolabs) for 30 min at 37°C in the dark at a final concentration of 250 nM. This TMR-probed suspension was centrifuged (12,000 x g, 3 min), then the cell sediment washed with 1 mL of PBS, suspended again in 1mL of PBS and labeled with Mitotracker Green (MTG, Thermofischer) for 1 min at room temperature at a final concentration of 1µg/mL. Cells were then washed three times in PBS and suspended in 50µL to 200 µL PBS, depending on the concentration of sporulating cells. For phase contrast and fluorescence microscopy, a 3 µL volume of the labeled cells suspension was applied onto 1.7% agarose coated glass slides and observed with an epifluorescence microscope (BX-61; Olympus) equipped with an Orca Flash 4.0 LT camera (Hamamatsu). Images were acquired using the CellSens Olympus software and micrographs were processed using ImageJ software. For quantification of the subcellular localization of SNAP-fusions, a minimum of 150 cells were counted and the different patterns identified were randomly examined and scored. Super Resolution Structured Illumination Microscopy (SR-SIM) images were acquired using an Elyra PS.1 Microscope (Zeiss) equipped with a Plan-Apochromat 63x/1.4 oil DIC M27 objective and a Pco. edge 5.5 camera, using 488 nm (100mW) or 561 nm (100 mW) laser lines, at 5–20% of total potency (27). The grid periods used were 23 mm, 28 mm or 34 mm for acquisitions with the 488 nm or 561 nm lasers respectively. For each SR-SIM acquisition, the corresponding grating was shifted and rotated five times, giving a total of 25 acquired frames. Final SR-SIM images were reconstructed using the ZEN software (black edition, 2012, version8, 1, 0, 484; Zeiss), using synthetic, channel-specific Optical Transfer Functions (OTFs). At least 100 sporangia/ 30 spores produced at 20°C were examined at each sampling times. The distance between cap and the forespore membranes was evaluated on at least 15 sporangia. All the kinetics were replicated at least two times using conventional fluorescence microscopy before being imaged once by SR-SIM.

### Transmission electron microscopy

Sporulating cells were centrifuged at 3,500 g during 5 min and fixed for 1h at room temperature and overnight at 4°C with 5% glutaraldehyde (v/v) in a 0.1 M sodium cacodylate buffer (pH 7.2) containing 1 mg. ml-1 ruthenium red. Cells suspended in 0.2 M sodium cacodylate were washed by three successive centrifugations (5 min at 3,500 g) and supernatant elimination and post-fixed for 1 h at room temperature with 2% osmium tetroxide. Cells were observed by transmission electron microscopy (TEM; Hitachi HT7800). A minimum of 30 sporangia were examined at the different times of sampling. The distance between the cap and the forespore membranes was scored on at least 15 sporangia.

## ACKNOWLEDGMENTS

Authors thank Dr Mariana Pinho for giving an access to the SR-SIM microscope, Pedro Matos for very efficient training of image acquisition and analysis software and Bénédicte Doublet for validation tests of *B. cereus* anti-CotE antibodies. Authors also thank Pr Anne Moir for gift of *exsY* and *cotY* mutant strains. This work is a partial fulfilment of AL Ph.D. thesis funded by INRAE and PACA Region and partly supported by a grant of MICA division and a Perdiguier grant of Avignon University. This work was also supported by FCT, “Fundaçã o para a Ciência e a Tecnologia”, Portugal through grant PEst-OE/EQB/LA0004/2011 to AOH and by program IF (IF/00268/2013/CP1173/CT0006) to MS. This work was also financially supported by project LISBOA-01-0145-FEDER-007660 (“Microbiologia Molecular, Estrutural e Celular”) funded by FEDER funds through COMPETE2020 – “Programa Operacional Competitividade e Internacionalizaçã o”, and by project PPBI - Portuguese Platform of BioImaging (PPBI-POCI-01-0145-FEDER-022122) co-funded by national funds from OE - “Orçamento de Estado” and by european funds from FEDER - “Fundo Europeu de Desenvolvimento Regional”. This work also was supported by the microscopy facilities of the Platform 3A, funded by the European Regional Development Fund, the French Ministry of Research, Higher Education and Innovation, the PACA region, Vaucluse Departmental Council and Avignon Urban Community.

## SUPPLEMENTAL MATERIAL

**Supplemental Results and discussion**

**FIG S1 to S6**

**TABLE S1 and S2**

## SUPPLEMENTAL FIGURE LEGENDS

**FIG S1** Stages of ExsY-SNAP and SNAP-CotE localization during sporulation. Sporulating cells of *B. cereus* producing ExsY-SNAP (A) or SNAP-CotE (B) were labeled with the SNAP substrate TMR-Star and with the membrane dye MTG and imaged by SR-SIM microscopy. Cells representative of the different localization patterns seen (*a* to *h*, see Fig. 1) are presented. TMR panels correspond to the signal from the SNAP-fusion, MTG corresponds to the membrane signal. Pink arrows point to exosporium at the different steps of its formation. Scale bar, 0.5 µm. (C) The percentage of sporulating cells showing the indicated patterns at the different times of sampling is presented for CotY-SNAP, ExsY-SNAP and SNAP-CotE.

**FIG S2** Fusion of the SNAP tag to the C-terminus of CotE affects separation of the cap. (A) Sporulating WT cells of *B. cereus* producing SNAP-CotE were labeled with the TMR-Star substrate and the MTG membrane dye and imaged by SR-SIM microscopy. The blue arrow points to a layer formed by CotE-SNAP distinct from the exosporium layer (pink arrows). This layer was not observed with the other SNAP-fusions tested (Fig. 1, S2A-B). Scale bar, 0.5 µm. Thin-sections of a representative CotE-SNAP-producing sporulating cell (B) after engulfment completion showing a prominent cap at the MCP forespore pole (black arrow) Scale bar, 200 nm; (C) following coat formation, the exosporium appeared to have a a normal structure. Scale bar, 500 nm. Ex, exosporium; Ct, coat.

**FIG S3** Absence of CotE impairs cap formation and leads to a modified pattern of coat deposition. Thin-sections of sporulating *B. cereus* WT ATCC 14579 (A to D) or *cotE* (E to H) cells, collected after hour 24 (A-B), 28 (C-D), 31 (E) or 48 (F-H) of sporulation at 20°C and observed by TEM. (A) a forespore presenting a cap at the MCP; (B) a cap separated from the forespore surface; (C) coat deposition on the longitudinal side or (D) coat deposition on the longitudinal sides and at the MCD forespore pole; (E-H) forespore of a *cotE* mutant after engulfment completion (E) without any visible coat or cortex (F) with a visible cortex and without any cap or exosporium material, in contrast with the WT at a similar stage of development (A-C); (G) a *cotE* forespore with altered coat assembly or (H) with large accumulation of exosporium material in the mother cell cytoplasm. FM, forespore membranes; Ex, exosporium material (cap in A-B-C); Cx, cortex; Ct, coat. Scale bar, 200nm.

**FIG S4** Localization of SNAP-CotE during sporulation of the *exsY* mutant. Sporulating cells of an *exsY* mutant producing SNAP-CotE were collected at the indicated times and analyzed by fluorescence microscopy. The *a*-*g* patterns are similar to the ones described for the WT producing SNAP-CotE (Fig. 1 and Fig. S1B). Numbers on the right indicate percentage of cells identified in each pattern (see Fig. 1 and main text). ND, the fluorescence signal from SNAP-CotE is not detected. The yellow arrow points to the cap structure typically observed in *exsY* spores. Scale bar, 1 µm. Coat/exosporium extracts were prepared from spores of WT ATCC 10876, the congenic *exsY* and *cotY* mutants (B) and derivative strains complemented through the ectopic production of ExsY-SNAP or CotY-SNAP (C) separated by SDS-PAGE and subject to immunoblotting with anti-CotE (bottom panels). The Coomassie-stained gels are shown as a loading control (top panels). One red asterisk indicates the CotE monomer, two red asterisks indicate a possible dimer, blue asterisks point to possibly cleaved forms; green asterisks point to non-specific bands (see also Fig. S6). In B and C, the position of molecular weight markers (MW, in kDa) is shown on the left side of the panels.

**FIG S5** CotY-SNAP, ExsY-SNAP and SNAP (the latter produced under the control of *cotE* promoter, p*cotE*-SNAP) accumulate in cell extracts throughout *B. cereus* sporulation. (A) WT cells with the indicated SNAP-fusions were harvested at the indicated times during sporulation. Extracts were prepared and used in pull-down assays and immunoblot analysis with anti-SNAP antibodies. An additional “spores” lane corresponds to pull-down done with extracts prepared from spores of the CotY-SNAP- and ExsY-SNAP-producing strains. One red asterisk indicates the indicated SNAP fusions; two or three red asterisks present potential multimeric forms of the SNAP-fusion proteins. Blue asterisk indicates cleaved forms. The position of molecular weight markers (MW, in kDa) is shown on the left side of the panels. (B) Cells expressing p*cotE*-SNAP were collected at hour 24 of sporulation, labeled with TMR-Star and MTG and imaged by phase contrast and fluorescence microscopy. The SNAP was detected dispersed throughout the mother cell cytoplasm. Scale bar, 1µm.

**FIG S6** CotE forms complexes with CotY in the cap of the *exsY* mutant. (A) Cells of an *exsY* mutant producing CotY-SNAP were harvested at hour 0, 24, 28, 48, and 72 of sporulation and whole cell extracts were submitted to pull-down and immunoblot analysis with anti-CotE and anti-SNAP antibodies. One red asterisks indicates CotE or a CotY-SNAP monomer; two red asterisks indicate potential multimers; blue asterisks point to possible products of proteolysis; green asterisks point to non-specific bands that react with the anti-CotE antibody on vegetative cell extracts. The position of molecular weight markers (MW, in kDa) is shown on the left side of the panels. (B) CotY-SNAP localization in the *exsY* mutant at hour 28, 48 and 72 of sporulation at 20°C. The numbers on the right of the panels indicate the percentage of cells identified in each pattern (see Fig. 1 and the main text). The panels show CotY-SNAP signals blocked at the cap stage. Scale bar, 1µm.

